# Nuclear lamina strain states revealed by intermolecular force biosensor

**DOI:** 10.1101/2022.03.07.483300

**Authors:** Brooke E. Danielsson, Bobin George Abraham, Elina Mäntylä, Jolene I. Cabe, Carl R Mayer, Anna Rekonen, Frans Ek, Daniel E. Conway, Teemu O. Ihalainen

**Author notes:** Contributed equally (Bobin George Abraham, Elina Mäntylä).

## Abstract

Nuclear lamins have been considered to be an important structural element of the nucleus. The nuclear lamina is thought both to shield DNA from excessive mechanical forces and to transmit mechanical forces onto the DNA. However, to date there is not yet a technical approach to directly measure mechanical forces on nuclear lamins at the protein level. To overcome this limitation, we developed a nanobody-based intermolecular tension FRET biosensor capable of measuring the mechanical strain of lamin filaments. Using this sensor, we were able to show that the nuclear lamina is subjected to significant force. These forces are dependent on nuclear volume, actomyosin contractility, functional LINC complex, chromatin condensation state, cell cycle, and EMT. Interestingly, large forces were also present on nucleoplasmic lamins, indicating that these lamins may also have an important mechanical role in the nucleus. Overall, we demonstrate that nanobody-based approach allows construction of novel force biosensors for mechanobiology studies.

## Introduction

Mechanical forces are important co-regulators of many physiological processes.^1^ In addition to mechanotransduction at the surface of the cell, the cytoskeleton also allows transmission of forces throughout the cell, including onto and within the nucleus.^2^ Thus, the nucleus has emerged as a putative mechanosensitive structure. At the nuclear envelope (NE), Linker of Nucleoskeleton and Cytoskeleton (LINC) complex, consisting of nesprin and SUN proteins, form the principal structure that connects the nucleus to the cytoskeleton.^3^ These connections enable a mechanotransmission pathway, where mechanical stress can be transduced bidirectionally between the cell surface and the nucleus via the cytoskeleton.^4^ Inside the nucleus LINC complex connects to nuclear lamina, a protein rich meshwork lining the inner nuclear membrane. The lamina is approximately 15 nm thick protein meshwork, formed mainly from flexible ~400 nm long A-type and B-type lamin filaments.^5,6^ Large parts of the chromatin are tethered to the nuclear lamina and this interaction has been shown to regulate gene expression.^5^ Especially A-type lamin proteins, lamin A/C, are also located throughout the nucleoplasm. These nucleoplasmic lamins bind to chromatin and have been indicated to regulate chromatin accessibility and spatial chromatin organization.^7^ Furthermore, mechanical stress has been shown to be transmitted deep into the nucleus and affect nuclear substructures, and even local chromatin organization and transcription in lamin A/C dependent manner.^2,8^ Thus, nuclear lamina forms a physical interface between chromatin and cytoplasm, and this interface is exposed to different mechanical cues.

*In vitro* experiments of purified nuclear lamins have shown that these proteins are able to withstand large mechanical forces, exhibit deformation under mechanical loading, and show strain-stiffening behavior.^9^ Inside the cells the amount of A-type lamins depends on cell substrate rigidity and nuclear lamina organization is affected by cellular contractility.^10,11^ Currently, there are no methods to directly detect changes in lamina strain state and this precludes studies on the effect of lamin strain-state in mechanotransduction between cytoskeleton and nucleus and its downstream implications. To study the mechanical loading of lamin filaments *in vivo* with subcellular resolution, we sought to develop a biosensor for lamin A/C. Prior force biosensor design strategies consisted of chimeric proteins in which a FRET pair separated by a force-sensitive peptide was inserted in the middle of the protein.^12^ These intramolecular force sensors have been successfully developed for proteins within focal adhesions,^13–15^ cell-cell adhesions,^16–20^ and the nuclear LINC complex.^21–23^ However, concerns remain regarding how internal insertion of a large FRET-force module may affect the biological functions of the protein. This may be especially important in the context of filamentous proteins, such as the nuclear lamins, where an altered protein size and structure may impair the assembly of filamentous structures.

Here we introduce a new concept for cellular mechanobiology studies, a nanobody-based intermolecular strain sensor. In this concept, instead of inserting a FRET-force module into the protein directly (intramolecular), we used nanobodies which bind to two proteins of interest (intermolecular). Our intermolecular strain sensor measures mechanical deformations between two proteins, in contrast to intramolecular force sensors which measure mechanical tension across a single protein. Our Lamin A/C strain sensor (Lamin-SS) consists of an existing FRET module (TsMod)^13^ is flanked on each side with nanobodies targeted against Lamin A/C.^24^ This indirect sensor design enables indirect tagging of endogenous proteins of interest without significantly impairing assembly, expression levels or cellular localization of these proteins. With this notion, we designed Lamin A/C strain sensor (Lamin-SS) using nanobodies targeted against Lamin A/C. Expressing this sensor in Madin-Darby canine kidney (MDCK) cells we demonstrate that this sensor exhibits an inverse FRET-force relationship. To validate the sensor, we used osmotic shock -induced nuclear shrinking and actomyosin contractility inhibitors. Using the sensor we are able to show that changes in chromatin condensation have a significant impact on lamin A/C force. Furthermore, our findings also show that the sensor experiences similar levels of tension in the nucleoplasm as compared to the nuclear lamina. The ability of nucleoplasmic lamin A/C to impart tensile forces was further supported by a second nanobody-based strain sensor which showed tensile forces between histone H2A-H2B and lamin A/C (Lamin-histone-SS) demonstrating also the versatility of the nanobody-based strain sensors. This technical advancement provides significant insights into nuclear mechanics, by providing the first direct measurements of nuclear lamina forces at the protein level. Additionally, the new sensor design demonstrates the potential for nanobody-based biosensors to be further utilized to measure mechanical forces between proteins.

## Results

### Nanobody-based lamin A/C strain sensor measures mechanical force exerted by nuclear lamins

We developed a lamin A/C strain sensor (Lamin-SS) that consists of an existing FRET-force module, known as TSmod^13^ with N- and C-terminal lamin A/C nanobodies^24^ (Fig. 1a). TSmod is a well-established FRET-force module which has an inverse FRET-force relationship. A force-insensitive truncated control sensor (Lamin-TM), containing only an N-terminal lamin A/C nanobody, was also developed (Fig. 1a). Next, we generated MDCK cell lines stably expressing Lamin-SS or Lamin-TM. The fluorescence of both sensors was strongly correlated to lamin A/C immunostaining (Fig. 1b), indicating that the sensors have strong localization to the nuclear lamina.

**Figure 1.**
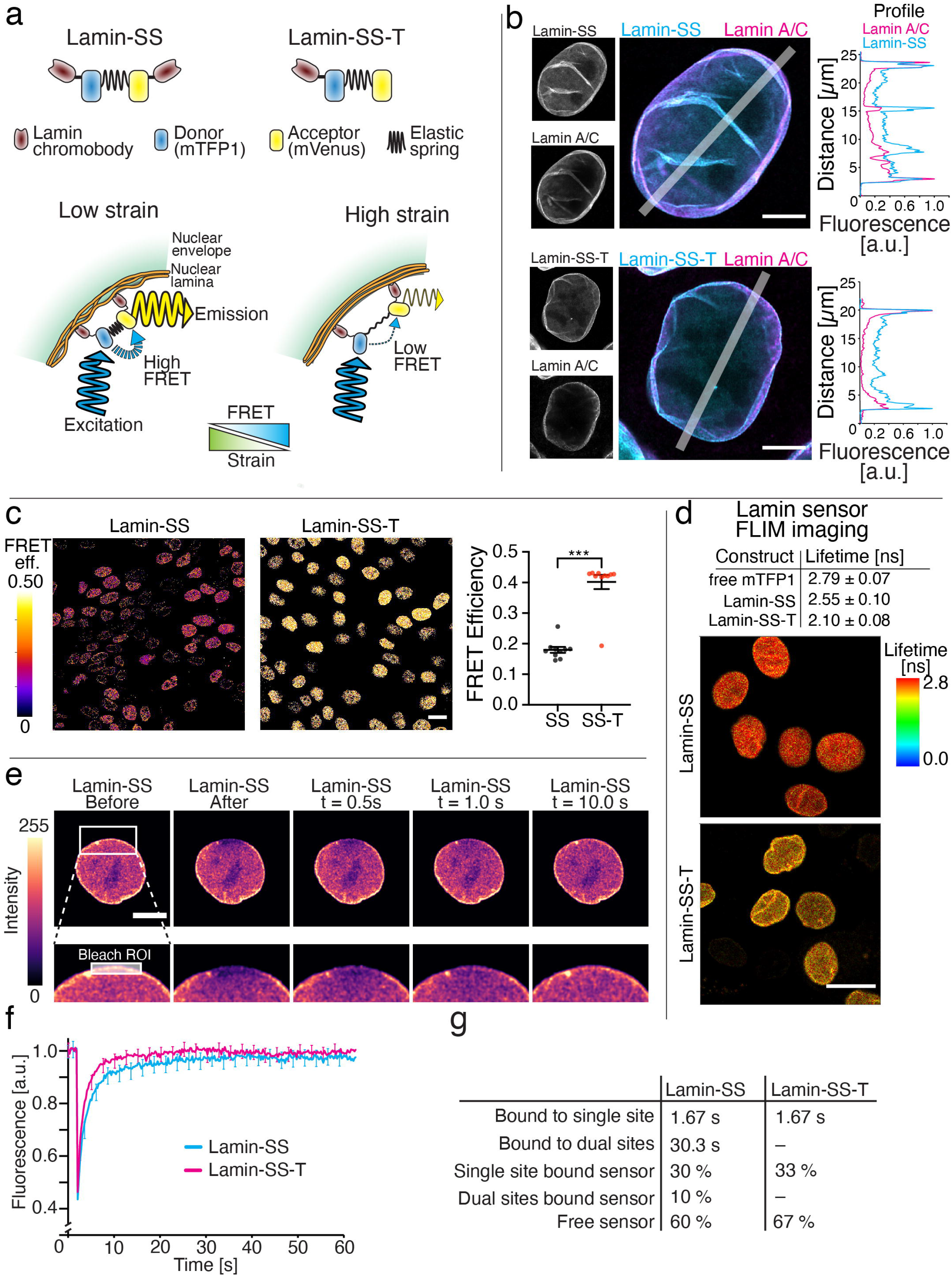
Development and characterization of the FRET based lamin A/C strain sensor. **a**, Schematic representation of the FRET-based lamin A/C strain sensor (Lamin-SS), truncated control sensor (Lamin-TM) and the working mechanism of the strain sensing. **b**, Confocal microscopy images of immunolabeled lamin A/C together with Lamin-SS or Lamin-TM and corresponding fluorescence line-profiles. Scale bars, 5 μm. **c**, FRET efficiency images and quantified FRET efficiency of Lamin-SS and Lamin-TM sensors (n=10 fields, 3 replicates). Scale bar 20 μm. Unpaired Student′s t-tests ((***) p<0.0001, t = 8.8, df = 18, p=0.0148). **d**, Donor fluorescence lifetimes of free donor (mTFP1), Lamin-SS and Lamin-TM along with FLIM images of Lamin-SS and Lamin-TM expressing cells (n=36, n=43 and n=42 cells, 2 replicates). Scale bar 20 μm. **e**, FRAP experiment with Lamin-SS expressing cell. Bleached region of interest (ROI) is marked in the blow-up image. Scale bar 5 μm. **f**, Quantified and normalized fluorescence recoveries (mean ± STDEV) of Lamin-SS and Lamin-TM indicating difference in the recovery dynamics (n=19 and n=18 cells, respectively, from 2 replicates). **g**, Lamin-SS and Lamin-TM binding times and corresponding fractions based on the simulated recoveries.

We measured FRET of each sensor using spectral-based FRET measurements. Lamin-SS exhibited reduced FRET as compared to Lamin-TM (Fig. 1c) indicating an increased distance between the FRET pair (assumed to be due to tension across the elastic peptide in TSmod). Lamin-SS had a median FRET efficiency of 17 % (mean 0.18 ± 0.01 SEM) compared to Lamin-TM with 40 % (mean 0.40 ± 0.02 SEM). As an additional control, we further confirmed these FRET changes using fluorescence-lifetime imaging microscopy (FLIM), which also showed reduced FRET (measured as increased lifetime) for Lamin-SS, as compared to Lamin-TM (Fig. 1d). To confirm that the FRET changes are dependent on A-type lamins, we constructed a MDCK cell line with CRISPR-Cas9 knockout of LMNA gene coding for A-type lamins. Loss of localization at the nuclear rim and higher FRET for Lamin-SS were observed in lamin A/C knockout cells (Supplemental Fig. 1), showing that the reduced FRET observed with Lamin-SS is dependent on A-type lamins. Apparent Lamin-SS FRET was 0.157 ± 0.007 in WT and 0.197 ± 0.005 in LMNA KO cells (mean ± SEM). Additionally, fluorescence recovery after photobleaching (FRAP) experiments and subsequent simulations of the sensor recovery behavior showed that both of the sensors, Lamin-SS and Lamin-TM, bind to nuclear lamina. The single nanobody based control sensor Lamin-TM had only a short binding time, but the dual nanobody sensor Lamin-SS showed substantially longer binding time to nuclear lamina (Fig. 1f, Fig. 1g, Supplemental Fig. 2 and Supplemental Fig. 3).

### Lamin A/C forces are dynamic, responding to changes in nuclear volume, actomyosin contractility, and nuclear-cytoskeletal connectivity

To further validate FRET-force responsiveness of Lamin-SS, we measured the FRET ratio during osmotic shock-induced nuclear shrinkage. In agreement with prior work,^25^ hyperosmotic shock reduced nuclear volume (Fig. 2a, 2b). Through paired cell analysis before and after sucrose-induced nuclear volume changes, we observed an increase in Lamin-SS FRET ratio, indicating less tensed sensor (Fig. 2c, 2d). Quantified Lamin-SS apparent FRET efficiency was 0.064 ± 0.001 in control conditions (MEM) and 0.114 ± 0.001 (mean ± SEM) in hyperosmotic conditions (MEM + 250 mM sucrose, 15 min).

**Figure 2.**
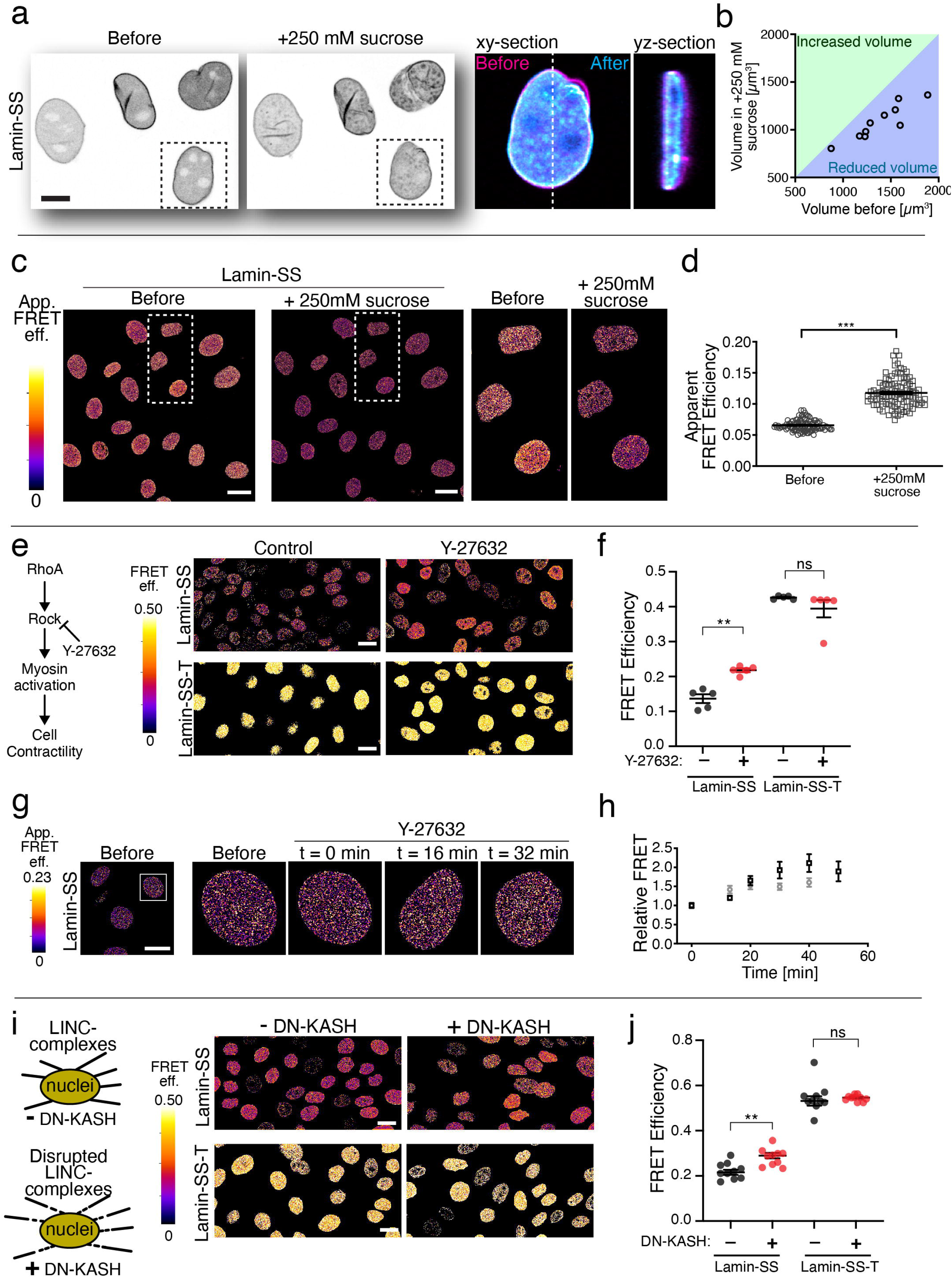
The effect of nuclear deformation, actomyosin contractility and nuclear force transduction on lamin A/C strain state. **a**, Lamin-SS expressing cells (left) were subjected to hyper-osmotic conditions by adding media containing 250 mM concentration of sucrose for 15 min (middle). Single cell blow-up indicated the change in nuclear morphology before and after the osmotic shock (right). Scale bar 10 μm. **b**, Scatterplot of quantified nuclei volumes indicating clear reduction of the nuclear volume. **c**, Apparent FRET efficiency images of osmotically stressed Lamin-SS expressing cells. Scale bar 10 μm. **d**, Quantified Lamin-SS apparent FRET efficiency. Plot represents the data (mean ± SEM) from a single measurement (total n=304 cells from 3 replicates). Wilcoxon matched-pairs test (*** p<0.0001). Scale bar 20 μm. **e**, FRET efficiency images of Lamin-SS and Lamin-TM after cell contractility inhibition (Y-27632, 50 μM, 1 h). **f**, Quantified FRET efficiency of Lamin-SS and Lamin-TM sensors after ROCK-inhibition (n= 10 fields, 3 replicates). Ordinary one-way ANOVA Tukey’s multiple comparisons (for Lamin-SS (**) p=0.005 and Lamin-TM (ns) p=0.4132). **g**, Lamin-SS apparent FRET efficiency imaging during ROCK-inhibition (Y-27632, 50 μM). **h**, Quantified relative change in Lamin-SS FRET ratio during ROCK-inhibition (n=152 cells from 2 replicates, black and grey). **i**, Disruption of LINC complexes by dominant-negative KASH (DN-KASH) expression (induction for 24h). FRET efficiency images of Lamin-SS and Lamin-TM after LINC disruption. **j**, Quantified FRET efficiency of Lamin-SS and Lamin-TM after LINC complex disruption (n=10 fields, 3 replicates). Ordinary one-way ANOVA Tukey’s multiple comparisons (for Lamin-SS (**) p=0.004 and Lamin-TM (ns) p=0.1722).

Next, we sought to determine if the actin cytoskeleton or actomyosin contractility contribute to forces on lamin A/C filaments. Treatment with actin fiber depolymerizing agent (cytochalasin D) reduced Lamin A/C forces, which was detected as increased FRET of the Lamin-SS (Supplemental Fig. 4). The apparent median FRET ratio increased from 4.3% before to 5.1% after the cytochalasin D treatment. Subsequently we studied the effect of myosin contractility on the nuclear lamina forces. When using Lamin-SS we observed that reduced actomyosin contractility via ROCK-pathway inhibitor (Y-27632) decreased lamin A/C forces (Fig. 2e, 2f). FRET efficiency of Lamin-SS was 14 % without treatment (mean 0.14 ± 0.01 SEM) and 22 % with Y-27632 (mean 0.22 ± 0.01 SEM). FRET efficiencies were 40 % (mean 0.43 ± 0.002 SEM) and 39 % (mean 0.39 ± 0.02 SEM) with Lamin-TM, respectively. Furthermore, the sensor was successfully used to temporally analyze lamin A/C force changes during actomyosin inhibition. The timelapse imaging data indicated that the FRET increased rapidly after the addition of the drug and 20 minutes of treatment showed significantly increased FRET (Fig. 2g, h). Thus, intact actin cytoskeleton and myosin contractility contribute to mechanical forces on lamin A/C filaments.

Because cytoskeletal tension is transduced to nuclear lamina in part by the nuclear LINC complex,^26^ we sought to understand the role of this structure for nuclear lamina forces. Disruption of the LINC complex using a dominant negative nesprin construct (DN-KASH)^26^ modestly reduced lamin A/C forces (Fig. 2i, j), further indicating the role of the cytoplasmic cytoskeleton for forces applied to the lamin A/C network. FRET efficiency of Lamin-SS was 22 % without DN-KASH expression (mean 0.16 ± 0.01 SEM) and 28 % (mean 0.19 ± 0.01 SEM) with DN-KASH expression. The efficiencies were 54% (mean 0.42 ± 0.001 SEM) and 53 % (mean 0.40 ± 0.01 SEM) with Lamin-TM, respectively. Taken together these data also demonstrate that lamin A/C forces are dynamic and they are affected by the cytoskeletal forces and intact LINC complexes. The data also demonstrates the dynamic responsiveness of the Lamin-SS sensor.

### Lamin A/C forces are affected by cell cycle, EMT, and chromatin condensation

Detecting a large heterogeneity in the quantified FRET values of Lamin-SS between individual cells, we hypothesized that cell-to-cell variations during the cell cycle could be affecting lamin A/C forces. When cells were arrested to early S-phase by treatment with DNA polymerase α inhibitor (aphidicolin), we detected increased Lamin-SS FRET (Fig. 3a), suggesting that lamin A/C forces are reduced by the cell cycle. Median FRET efficiency of Lamin-SS was 16 % without Aphidicolin (mean 0.16 ± 0.004 SEM) and 21 % (mean 0.21 ± 0.004 SEM) after Aphidicolin treatment. The efficiencies were 40% (mean 0.39 ± 0.02 SEM) and 39 % (mean 0.39 ± 0.02 SEM) with Lamin-TM, respectively. However, the cell-to-cell variation in force persisted indicating that additional factors beyond cell cycle are creating this heterogeneity in forces.

**Figure 3.**
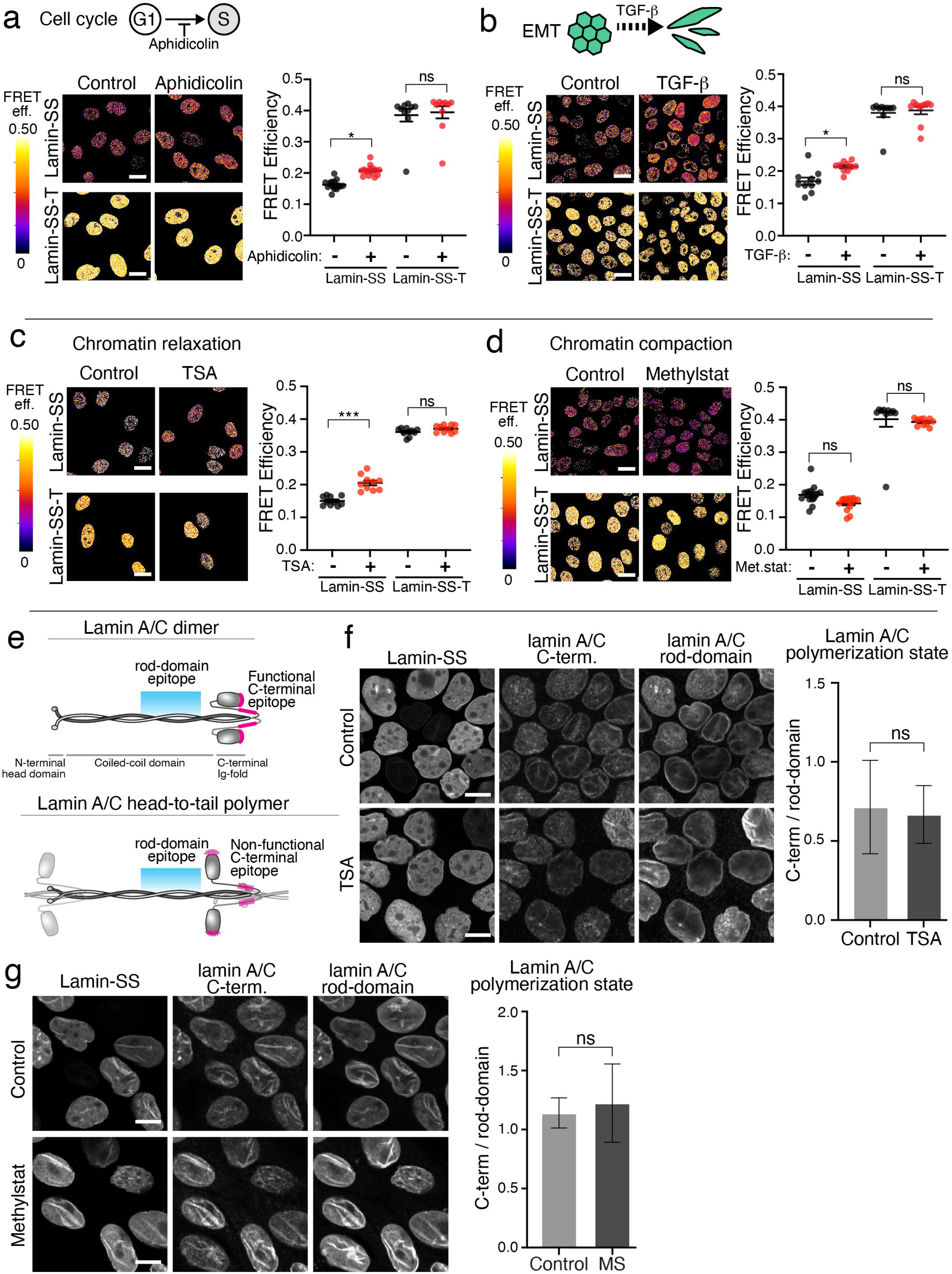
The effect of cell cycle, ETM and chromatin organization on lamin A/C strain. **a**, Cells synchronization to early S-phase (Aphidicolin, 3 μg/mL, 24 h). Images and quantification of Lamin-SS and Lamin-TM FRET efficiency after cell cycle synchronization (n= 10 fields, 3 replicates). Ordinary one-way ANOVA Tukey’s multiple comparisons (for Lamin-SS (*) p=0.0229 and Lamin-TM (ns) p=0.9619). Scale bars, 20 μm. **b**, EMT induction by growth factor treatment (TGF-β1, 2 ng/mL, 24 h). Images and quantification of Lamin-SS and Lamin-TM FRET efficiency after EMT induction (n=10 fields, 3 replicates). Ordinary one-way ANOVA Tukey’s multiple comparisons (for Lamin-SS (*) p=0.0311 and Lamin-TM (ns) p=0.9666). Scale bars, 20 μm. **c**, Chromatin relaxation by histone deacetylase inhibitor (TSA, 200 nM, 4 h). Images and quantification of Lamin-SS and Lamin-TM FRET efficiency after treatment (n=10 fields, 3 replicates). Ordinary one-way ANOVA Tukey’s multiple comparisons (for Lamin-SS (***) p<0.0001 and Lamin-TM (ns) p=0.4804. Scale bars, 20 μm. **d**, Chromatin condensation by histone demethylase inhibitor (methylstat, 2.5 μM, 48 h). Images and quantification of Lamin-SS and Lamin-TM FRET efficiency after treatment of the cells. Ordinary one-way ANOVA Tukey’s multiple comparisons (for Lamin-SS (ns) p=0.2282 and Lamin-TM (ns) p=0.9584). Scale bars, 20 μm. **e**, Localization of epitopes for the used lamin A/C antibodies. The C-terminal epitope accessibility depends on lamin filament organization. **f**, Imaging and quantification of lamina organization in TSA treated cells. Confocal microscopy maximum intensity projections of control (upper panels) and TSA-treated (600 nM, 4 h, lower panels) Lamin-SS expressing cells, immunolabeled against C-terminal and rod-domain epitopes of A-type lamins. Scale bars, 10 μm. Quantified fluorescence intensity ratio of lamin A/C labeling in control and TSA-treated cells (n=291 and n=276 cells, respectively, from 3 biological replicates). Unpaired Student′s t-test ((ns) p=0.6, t=0.5, df=28). **g**, Imaging and quantification of lamina organization in methylstat treated cells. Confocal microscopy maximum intensity projections of control (upper panels) and methylstat-treated (2.5 μM, 48 h, lower panels) Lamin-SS expressing cells, immunolabeled against lamin A/C C-terminus and rod-domain epitopes. Scale bars, 10 μm. Quantified fluorescence intensity ratio of lamin A/C labeling (n=244 and n=251 cells, respectively, from 3 biological replicates). Unpaired Student′s t-test ((ns) p=0.4, t=0.9, df=28).

Induction of epithelial to mesenchymal transition (EMT) by using TGF-β also resulted in decreased lamin A/C forces (Fig. 3b). Median FRET efficiency of Lamin-SS was 17 % (mean 0.17 ± 0.01 SEM) without TGF-β1 and 22 % (mean 0.21 ± 0.004 SEM) after TGF-β1 treatment. The efficiencies were 39 % (mean 0.39 ± 0.01 SEM) and 39 % (mean 0.38 ± 0.01 SEM) with Lamin-TM, respectively. Prior studies have suggested that chromatin decondensation increases during EMT.^27^ Additionally, chromatin condensation changes have been shown to affect nuclear stiffness,^28,29^ and thus may also have the potential to regulate the mechanical state of the nuclear lamins. To directly modulate chromatin condensation, we used a histone deacetylase inhibitor (TSA) and a histone trimethyl demethylase inhibitor (methylstat) to decondense and condense chromatin, respectively. Cells treated with TSA to decondense chromatin had significantly decreased lamin A/C force (Fig. 3c). FRET efficiency of Lamin-SS was 15 % without TSA (mean 0.15 ± 0.004 SEM) and 21 % after TSA treatment (mean 0.21 ± 0.01 SEM). The efficiencies were 37 % (mean 0.37 ± 0.004 SEM) and 36 % (mean 0.36 ± 0.004 SEM) with Lamin-TM, respectively. The increased histone acetylation after TSA treatment was confirmed by significantly higher immunostaining intensity of acetylated H3K27 (Supplemental Fig. 5). We also investigated the possibility that the lamin A/C organization is affected by the TSA and subsequently affects the Lamin-SS FRET. Here we used antibodies which recognize two different epitopes in the lamin A/C. The C-terminal epitope labeling has been shown to depend on higher organization of lamin A/C filaments^11^ as the other antibody recognizes the rod-domain region of the lamin A/C (Fig. 3e). Ratiometric imaging by using the antibodies indicated that the TSA treatment did not alter the organization of A-type lamins (Fig. 3f). In control cells the antibody labeling ratio was 0.72 ± 0.30 and in the TSA treated cells 0.69 ± 0.18. In addition, the treatment did not affect Lamin-SS binding to the nuclear lamina and the binding dynamics unaltered as shown by the FRAP experiments (Supplemental Fig. 6). In comparison to TSA induced decondensation of chromatin, cells treated with methylstat to condense chromatin exhibited a small but non-significant increase in lamin A/C force (Fig. 3d). Median FRET efficiency of Lamin-SS was 18 % (mean 0.17 ± 0.01 SEM) without methylstat and 14.5% (mean 0.14 ± 0.01 SEM) after methylstat treatment. The efficiencies were 40 % (mean 0.40 ± 0.02 SEM) and 39 % (mean 0.39 ± 0.003 SEM) with Lamin-TM, respectively. Similarly, to TSA treatment, methylstat also did not affect A-type lamin organization (Fig. 3g), in control cells the antibody labeling ratio was 1.14 ± 0.13 and in the methylstat-treated cells 1.22 ± 0.33.

### Nucleoplasmic lamins also experience mechanical force

Intriguingly, we detected similar levels of FRET at the nuclear perimeter and in the nucleoplasm (Fig 1c, 1d). In addition, we quantified the Lamin-SS and Lamin-TM FRET in the nuclear rim and nucleoplasm by using FLIM (Fig. 4a). The data indicated that the lifetime and thus the FRET did not change between the nuclear perimeter and nucleoplasm (Fig. 4b). Together these data suggest that nucleoplasmic A-type lamins also experience significant forces. Nucleoplasmic lamins have been shown to be assembled and to interact with chromatin.^30^ To further examine nucleoplasmic lamins, as well as the potential for force transmission between nuclear lamins and chromatin, we developed an additional nanobody sensor, using a previously developed H2A-H2B nanobody,^31^ to measure mechanical tension between histone H2A-H2B and lamin A/C (Lamin-histone-SS) (Fig. 4c). This sensor exhibited a more predominant nucleoplasmic localization than Lamin SS and was highly correlated to H2A staining (Fig. 4d). The Lamin-histone-SS exhibited reduced FRET in comparison to a truncated sensor Lamin-histone-TM (Fig. 4e). Lamin-histone-SS had a median FRET efficiency of 17 % (mean 0.27 ± 0.02 SEM), compared to Lamin-histone-TM with 40 % (mean 0.41 ± 0.02 SEM). For further validation Lamin-histone-SS was examined in the MDCK LMNA knockout cell line (Fig. 4f). Lamin-histone-SS exhibited higher FRET (reduced force) in the knockout cell line, indicating that Lamin A/C proteins are necessary for tension across this sensor (Fig. 4f, 4g). Lamin-histone-SS apparent FRET was 0.191 ± 0.006 in WT and 0.222 ± 0.0086 in LMNA KO cells, and Lamin-histone-TM apparent FRET was 0.261 ± 0.004 in WT cells and 0.271 ± 0.003 in LMNA KO cells (mean ± SEM). Although lamins and histones are known to physically associate,^32^ the Lamin-histone-SS establishes that mechanical forces can be transduced between chromatin and A-type lamins. Taken together, these results with both the Lamin-SS and Lamin-histone-SS indicate that nucleoplasmic lamins experience and can transmit significant levels of mechanical force.

**Figure 4.**
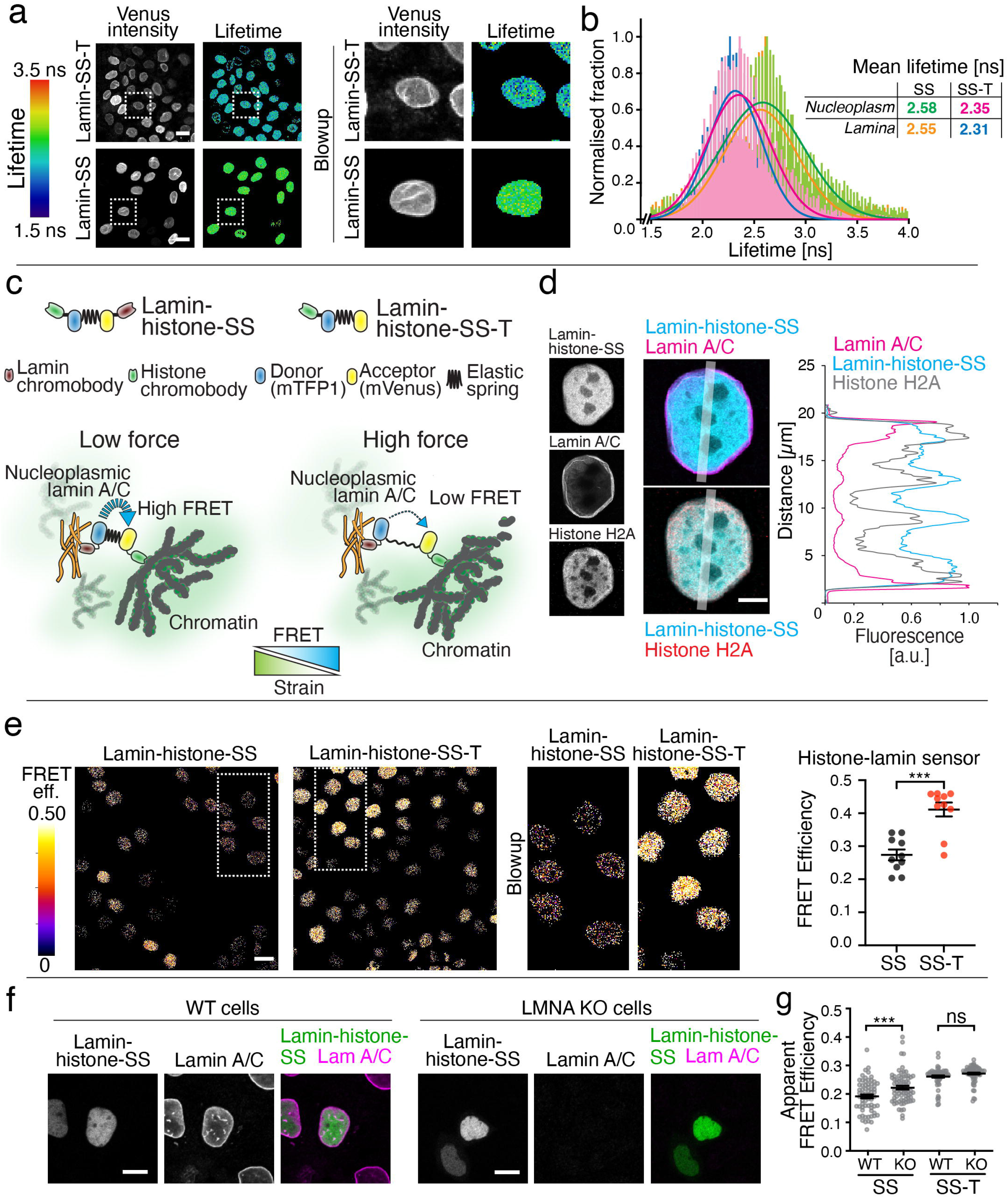
Mechanical strain in nucleoplasmic lamin A/C filaments. **a**, Acceptor (venus) intensity image together with donor (mTFP1) fluorescence lifetime microscopy images of Lamin-SS and Lamin-TM expressing cells. Scale bar 20 μm. Blowup images show equal distribution of lifetimes throughout the nucleus. **b**, Donor fluorescence lifetime histograms show highly similar lifetimes and thus FRET for nuclear rim and nucleoplasm, indicating similar strain in lamin A/C in the nuclear lamina and in the nuclear interior (n=26 and n=26 cells for Lamin-SS and Lamin-TM, respectively, 2 replicates). **c**, Schematic representation of the FRET based lamin A/C – histone H2A strain sensor (Lamin-histone-SS), truncated control sensor (Lamin-histone-TM) and the working mechanism of the force sensing between lamins and chromatin. **d**, Laser scanning confocal microscopy images (Airy-scan, single sections) of immunolabeled lamin A/C, histone H2A and the expressed Lamin-histone-SS sensor along with corresponding fluorescence line-profiles. Scale bar 5 μm. **e**, FRET efficiency images and quantified FRET efficiency of Lamin-histone-SS and Lamin-histone-TM sensors. The plots represent the median ± SEM of individual image fields (total n = 10 from 3 replicates). Scale bar 20 μm. Unpaired Student′s t-test ((***) p<0.0001, t=5.14, df=18). **f**, WT and LMNA KO cells transiently transfected with Lamin-histone-SS. Scale bars 10 μm. **g**, Quantified apparent FRET efficiency (mean ± SEM) of Lamin-histone-SS (n=67 and n=72, respectively, 2 replicates) and Lamin-histone-TM (n=67 and n= 78 in 2 replicates) in WT and LMNA KO cells. Ordinary one-way ANOVA Tukey’s multiple comparisons (for Lamin-histone-SS (***) p=0.0003 and Lamin-histone-TM (ns) p=0.4756)

## Discussion

Forces and mechanical stresses at the cellular and protein levels are difficult to quantify directly. Although a number of approaches have used externally applied force to study mechanotransmission of force onto and within the nucleus,^2,26^ more recently we and others have developed force-sensitive biosensors for nesprin proteins,^21–23^ as well as identified force-sensitive antibodies,^11^ which are technical approaches that can be used to characterize nuclear forces without the need for externally applied forces. In this work we established a new nanobody-based FRET-strain sensor concept, which enabled mechanical studies of the protein dense and filamentous nuclear lamina. Using this biosensor, we demonstrate the lamin A/C network experiences significant mechanical force, which is regulated by actin, myosin, and nuclear-cytoskeletal connections (Fig. 2). Additionally, we show that there exists a positive correlation between chromatin condensation and lamin A/C forces (Fig. 3). Intriguingly the nucleoplasmic lamin A/C forces were of a similar level to the forces of lamin A/C at the NE (Fig. 4).

There are a large number of prior studies demonstrating that the nucleus can experience and sustain mechanical loading. Physical perturbations of cells and isolated nuclei have shown that nuclear lamins deform under mechanical loading.^33,34^ Nuclear lamina is also a mechanosensitive structure and its organization changes according to cellular tensile state or cell substrate.^10,11^ Furthermore, forces are transduced through the nuclear lamina and can affect chromatin organization.^2^ Mechanical integrity of the nucleus has also been shown to depend on A-type lamins.^35^ In line with this, loss and mutant forms of lamin A/C lead to force-induced nuclear rupture and DNA damage.^34,36,37^ These studies indicate that nuclear lamina experiences and responses to mechanical cues. Our newly developed strain sensor adds to this prior work by demonstrating that the lamin A/C network is subject to constitutive mechanical tension in adherent cells. This kind of direct measurement of tensile state or mechanical strain of the lamina has not been previously possible.

In addition to localizing to the nuclear periphery (at the NE), A-type lamins are present throughout the nucleoplasm .^5^ Work by Roland Foisner and colleagues have shown that nucleoplasmic lamins associate with LAP2α and may have important roles in regulation of chromatin.^7^ Loss of LAP2α has been shown to decrease nucleoplasmic lamin mobility, resulting in a potentially more assembled filamentous nucleoplasmic lamin structure and decreased chromatin motion.^30^ Our observation of significant mechanical forces on nucleoplasmic A-type lamins (Fig. 4) further supports the idea that these lamins provide important structural and mechanical properties to the nucleus. Additionally, our histone-lamin sensor shows that nucleoplasmic lamins, as opposed to lamins at the periphery, are the primary lamin component mechanically interacting with chromatin (Fig. 4).

Our FRAP experiments demonstrated simultaneous, dual binding of both nanobodies in Lamin-SS to lamin A/C filaments (Fig. 1). Since the nanobody recognizes a single epitope in A-type lamins, it is possible that each nanobody in the sensor is binding to two lamin A/C proteins in a single filament. Alternatively, the sensor may also simultaneously bind to two lamin A/C proteins that are located in separate yet closely associated filaments. Lamin protein dimers form partly staggered head-to-tail polymers with repetitive features every 40 nm.^6^ Lamin filaments are assembled from two laterally interacting head-to-tail polymers and show 20 nm periodicity.^6^ Since we estimate that Lamin-SS may be up to 40 nm in length (when fully extended), either or both binding scenarios are possible.

A further hindrance in our understanding of how Lamin-SS is interacting with the Lamin A/C network is that the epitope of the lamin A/C nanobody has not yet been identified. We also note that we previously reported that mechanical forces can regulate lamin A/C epitope accessibility.^11^ Although our control experiments (Fig. 3) did not show major changes in accessibility of other lamin A/C epitopes with TSA and methystat treatments, it may be possible that the Lamin-SS epitopes are also influenced by the physical arrangements of the nuclear lamina, in addition to nuclear strain. However, our FRAP data indicates that TSA treatment does not influence Lamin-SS binding to the lamina (Supplemental figure 6), further arguing against force-induced changes in binding of Lamin-SS.

We also considered the possibility that changes in lamin A/C structure could affect the density of the sensor. Extremely close packing of multiple sensors can lead to intermolecular FRET, FRET occurring between neighboring sensors when sensors are closely packed together.^38^ We note that the truncated Lamin-TM did not exhibit FRET changes in response to experimental treatments (Fig. 2f, 2j, 3a, 3b, 3c, 3d, 4f). Because Lamin-TM should also be subjected to similar changes in sensor densities and packing as Lamin-SS during these treatments, we assume that treatment-induced changes in Lamin-SS FRET are occurring not from changes in intermolecular FRET, but instead changes in tensile forces across a single sensor.

Through our FRAP analysis of binding, we observed that the turnover rates of the Lamin-SS (Fig. 1) are much faster than the longer turnover rates of Lamin A.^39,40^ Thus, the reduced FRET of Lamin-SS, which has a dual-binding time of approximately 30 seconds, is due to dynamics of the lamin network and the turnover of lamins is not affecting the process. Our FRAP analysis indicates only 10 % of Lamin-SS exists with both nanobodies simultaneously bound (Fig 1g), which is surprising given the large decrease in FRET observed with this sensor as compared to Lamin-TM. One possibility is there is a delay in the relaxation of the sensor after unbinding.

While Lamin-SS sensors may bind, unbind, and re-bind rapidly, the extension of the flagelliform peptide may persist over a longer timescale, remaining extended even when the sensor is in a single or unbound state. Similarly, it is possible that mechanical forces applied across the sensor causes one or more of the fluorescent proteins to unfold,^41^ which would result in decreased fluorescence of the protein^42^ and therefore reduced FRET. If true, the refolding of the fluorescent proteins would likely occur on a longer timescale, resulting in decreased FRET persisting in the single or unbound states. If there is delayed refolding or relaxation of the sensor this raises the possibility that there would exist a temporal lag in Lamin-SS detection of force relaxation. We note that the sensor was readily able to detect lamin A/C relaxation after 15 minutes of osmotic nuclear shrinkage (Fig. 2c) and approximately 20 minutes of Y-27632 treatment (Fig. 2g), indicating that any temporal lag is modest and that force-induced changes in Lamin-SS are reversible.

While our approach might be considered to be the first use of TSmod in an intermolecular strain/force sensor (between two proteins), we note the prior work of Alex Dunn and colleagues in which TSmod was also used similarly in an intermolecular manner to measure forces exerted between integrins and the extracellular matrix.^43^ An important benefit to our nanobody-based intermolecular sensor is that it remains completely genetically encodable, allowing for cellular expression of the sensor. Furthermore, the implementation of nanobodies with TSmod enables this approach to be adapted to other proteins for which there are existing nanobodies, such as actin^44^ and vimentin.^45^ Additionally, existing nanobody epitope tags (C-Tag, Spot-tag, and ALFA-tag),^46^ as well as anti-GFP and anti-RFP nanobodies,^47^ could enable the development of nanobody strain sensors for virtually all proteins.

This technical advancement provides significant insight into nuclear mechanics, by providing the first direct protein-level measurements of nuclear lamina strain. Lamin A/C strain, presumably the result of tensile and compressive mechanical forces, is dynamic and influenced in both an outside-in (actomyosin, LINC complex) and inside-out (chromatin) manner. Additionally, we show that intranuclear lamins also experience significant levels of strain, providing additional evidence that nucleoplasmic lamins are an important structural element of the nucleus. This work demonstrates the potential for nanobody-based biosensors to be further utilized to measure mechanical forces between homotypic and heterotypic protein associations.

## Methods

### Sensor Design

The sensor to measure mechanical forces on the nuclear lamins was designed using an existing lamin A nanobody. The nanobody was previously developed by Rothbauer et al^24^ and is currently commercially distributed as a “chromobody”, consisting of the nanobody tagged with GFP, by Chromotek (Planegg-Martinsried, Germany). The sensor is designed such that an existing FRET-force biosensor, known as TSmod^13^ is flanked on either side by the lamin A nanobody V_H_H sequence (Fig. 1a). To ensure nuclear localization of the protein a c-myc NLS was inserted between each nanobody and TSmod. Additionally, the C-terminal lamin A nanobody was designed by using the reverse sequence of the V_HH_ for orientation of the nanobody outwards from TSmod. The entire sequence of the sensor was synthetically cloned by GeneArt (Thermo Fisher Scientific) into pcDNA 3.3. The plasmid is available through Addgene (plasmid# 178641). A control force-insensitive lamin sensor, consisting of only one nanobody attached to TSmod was also developed and is available through Addgene (plasmid# 178642).

A second sensor was developed designed to measure forces between the nuclear lamina and histones. This sensor consists of a nanobody which binds to the Histone H2A-H2B heterodimer that was previously developed by Jullien et al^31^ and is also commercially distributed as a “chromobody” by Chromotek. The sensor was designed similarly to the lamin sensor, with the N-terminal lamin A sensor being replaced with the histone nanobody (histone nanobody-TSmod-reverse lamin A nanobody). This sensor was also synthetically cloned by GeneArt and is available through Addgene (plasmid# 178643). A control force-insensitive histone sensor, consisting of only the histone nanobody attached to TSmod was also developed and is available through Addgene (plasmid# 178644).

### Cells

Madin-Darby canine kidney cells (MDCK II) were used in all studies and maintained in high glucose DMEM (Thermo Fisher Scientific) supplemented with 10% fetal bovine serum (Thermo Fisher Scientific) and 1% penicillin/streptomycin (Thermo Fisher Scientific) under standard cell culture conditions. To generate stable cell lines, MDCKs were transfected with the TSmod and selected using G418. For the DN-KASH experiments, DN-KASH inducible Lamin-SS cells and DN KASH inducible Lamin-TM cells were made into stable cell lines. To generate a system for doxycycline-inducible DN-KASH Lamin-SS cells and doxycycline-inducible DN-KASH Lamin-TM cells, the previously established doxycycline-inducible DN-KASH MDCK cells^48^ were electroporated with Lamin-SS pcDNA and Lamin-SST pcDNA separately. Cells expressing both DN-KASH and Lamin-SS/TM were extracted with cloning rings and were clonally expanded.

### Establishment of LMNA knockout with CRISPR/Cas9

To generate a pre-LMNA knockout (KO) MDCK II cell line with CRISPR/Cas9, single guide RNAs (sgRNAs) were custom made from Invitrogen backbone from their LentiArray™ Human CRISPR Library and designed against LMNA1 gene N-terminus in CanFam 3.1 reference genome (https://www.ncbi.nlm.nih.gov/nuccore/NM_001287151.1, GeneID: 480124) with an online guide design tool. LMNA target sequence: atggagac cccgtcccag cggcgcgcca cccgtagcgg ggcgcaggcc agctccaccc cgctgtcgcc cacccgcatc acccggctgc aggagaagga ggacctgcag gagctcaatg accgcctggc ggtctacatc gaccgtgtgc gctctctgga gacggagaac gcggggctgc gccttcgcat caccgagtcg. The sgRNA_LMNA_N1 nucleotide sequence was CACGGTCGATGTAGACCGCC (on-target locus chr7:-41719582). For expression, the sgRNA_LMNA_N1 (300 ng) and pCDNA3.1-dCas9-2xNLS-EGFP (gift from Eugene Yeo, #74710, Addgene; http://n2t.net/addgene:74710) were transfected by using the Neon™ electroporation system (1650 V, 20 ms, 1 pulse; Thermo Fisher Scientific) followed by selection of GFP-positive cells with G418 (0.75 mg/mL, Merck) and FACS sorting (BD FACSAria Fusion, BD Biosciences).^49^ The KO cells was verified via Immunoblotting and further confirmed by immunostainings.

### Immunoblotting

For immunoblotting, MDCK II and MDCK LMNA KO cells were washed with PBS and lysed in buffer containing 50 mM Tris-Cl pH 7.5, 1% Triton X-100, 1 mM EDTA, 150 mM NaCl, 50 mM NaF, 10% glycerol, 1 mM phenylmethanesulfonyl fluoride, 8.3 μg/ml aprotinin, 2 mM vanadate, and 4.2 μg/ml pepstatin. The lysates were centrifuged and used for SDS-PAGE with Mini-PROTEAN^®^ TGX™ Precast gel (Bio-rad Finland OY). Following transfer to Amersham™ Protran^®^ nitrocellulose blotting membrane, the immunoblots were blocked with 2% BSA and incubated with primary antibodies: Anti-Lamin A/C antibody (E-1) (sc-376248, Santa Cruz Biotechnology) and Anti-Actin antibody, clone C4 (MAB1501R, Merk-Millipore). The primary antibodies were detected using a mixture of goat-anti-mouse (DyLight 800) and goat-anti-rabbit (DyLight 680) (both from Thermo Scientific) secondary antibodies and the signal was read using Odyssey CLx (LI-COR).

### Drug Treatments

For actin depolymerization studies, Cytochalasin D (cat # 11330, Caymen Chemical) was used at 10 μg/mL for 1 h. To inhibit Rho A kinase, 50 μM Y-27632 (cat #72302, Stem Cell Technologies) was used for 1 hour prior to FRET imaging to reduce myosin activity. For EMT induction, recombinant human TGF-β1 (R&D systems) was used to induce EMT at a concentration of 2 ng/mL for 24 h. Modifications in DNA ultrastructure were done to condense or decondense chromatin with the use of 600 nM trichostatin A (TSA) for 4 h (Cayman Chemical Company), to increase euchromatin, and 2.5 μM methylstat (Sigma Aldrich) for 48 h, to increase heterochromatin. For the cell cycle synchronization assay, Aphidicolin (cat #57-361, Thermo Fisher Scientific) was used to block the cells in early S-phase, at 3 μg/mL for 24 h.

### Immunofluorescence Staining of Lamins and Histones

For fixed-cell experiments, cells were washed with PBS and fixed for 10 min at room temperature with 4% paraformaldehyde in PBS. After three washes with PBS, the cells were permeabilized for 10 min at room temperature with 0.2% Triton X-100 in PBS and blocked with 5% BSA for 1 h at room temperature. Cells were then incubated overnight at 4 °C or in room temperature with the primary Ab diluted in blocking solution. The following primary antibodies were used: anti-lamin A antibody (sc-7292, Santa Cruz Biotechnology), anti-mouse LA/C-C (131C3, ab8984, Abcam) anti-rabbit LA/C-rod (EP4520-16, ab133256, Abcam), or anti-Rabbit Histone H2A (cat #12349, Cell Signaling Technology). Three more washes with PBS were then followed by incubation with the secondary Ab (Alexa Fluor 647-conjugated donkey anti-mouse IgG; Thermo Fisher Scientific) for 45 min followed by three additional PBS washes. Samples were stained with Hoechst 33342 (Thermo Fisher Scientific) and mounted with ProLong Gold antifade mountant (P36930, Thermo Fisher Scientific).

### Fluorescence Microscopy of Histones and Nuclear Lamina Organization

Confluent non-treated or either trichostatin A (TSA, 4h, 600 nM) or methylstat (48h, 2.5 μM) – treated MDCK II wt, MDCK II-TS or MDCK II-truncated mutant cells were analyzed to ensure the drug treatments used in FRET-experiments did not affect the nuclear lamina organization. To detect changes in nuclear lamina organization, ratiometric fluorescence immunoassay was performed on MDCK II wt, MDCK II cells stably expressing Lamin-SS or MDCK II cells stably expressing Lamin-TM immunostained against either lamin A/C N-terminus (LA/C-N, E1, sc-376248, Santa Cruz Biotechnology, Texas, USA) and histone H3 lysine 27 acetylation (H3K27ac, ab4729, Abcam), or lamin A/C C-terminal (anti-mouse LAC/C-C, 131C3, ab8984, Abcam, Cambridge, UK) and lamin A/C rod domain (anti-rabbit LA/C-rod, EP4520-16, ab133256, Abcam) as described above. Imaging was done on a Nikon A1R+ laser scanning confocal mounted in Nikon Eclipse Ti2-E inverted microscope (Nikon Instruments, Tokyo, Japan). Nikon 60X/1.40 Apo DIC N2 oil immersion objective was used in the experiments. Solid state lasers with excitation wavelengths 488 nm, 561◻nm and 640 nm were used in excitation. The emissions were collected with 525/50, 540/30 and 595/50 bandpass filters, respectively. The laser intensities were adjusted to avoid photobleaching and the detector sensitivity was adjusted to optimize the image brightness and to avoid saturation. Laser powers and detector voltages were determined individually per treated antibody pair, and after the initial setting kept constant for each sample to allow ratiometric imaging and quantitative comparison of the fluorescence intensities within the drug-treated and non-treated control samples. The images were 1024×1024 pixels and the pixel size was 103.6 μm in x/y. The images were acquired without averaging and by first focusing on the bottom surface of the sample, where the position of the sample stage was set as ◻0=0. The fluorescence signal intensities from all emission channels were then collected from bottom to top as optical z-series with 200◻nm step size. The pinhole was set to 0.9 (physical pinhole size 34.76 μm). The analysis was done in ImageJ software by making maximum intensity projections from the acquired z-stacks, and by using the LA/C-rod channel to segment the nuclei which was then used as a mask to measure the maximum signal intensities for all channels. The mean intensities of the nuclei, the background, and the total images were determined. To detect changes in the lamin organization the nuclear lamin intensity ratio (LA/C-C:LA/C-rod) was calculated from the nuclear intensities of which the detector noise was subtracted and which were normalized against their background. Number of replicates, n=3 for all treatments. Unpaired Student′s t-test was used to test the statistical significance between treated and non-treated control samples. ns= non-significant, *p<0.05, **p<0.01, ***p<0.001.

### Super-resolution Airy-scan Imaging

Zeiss LSM 980 laser scanning confocal microscope with airyscan was used for fixed-cell experiments. The system was mounted on Axio Observer.Z1 microscope body and Plan-Apochromat 63X/1.4 oil immersion objective was used in the imaging. The sensor and the immunolabelled lamin A/C were excited with 488 nm and 639 nm lasers using MDS488/561/639 triple dicroic and the emission was collected with band-pass 495-560 nm and long-pass 650 nm filters. The image size was set to 1032 x 1032 pixels, with pixel size of 43 nm and optical section collected with 170 nm intervals. Scanning was bidirectional with 2 μs pixel dwell time and averaging of 4 was used. Data was analysed with ImageJ FIJI -distribution.

### Fluorescence Recovery After Photobleaching (FRAP)-Experiments

Zeiss LMS780 laser scanning confocal microscope in inverted Cell observer microscope body was used in the experiments. MDCK cells stably expressing Lamin-SS or Lamin-TM were seeded on collagen-I -coated (50 μg/mL in PBS, 45 min in RT) high performance coverslips (Zeiss, #474030-9020-000) 1 d before the experiments. Prior imaging, the coverslips were mounted on imaging chamber (Aireka Cells, #SC15022, Aireka Scientific, HK, China) and placed in the microscope incubator (37 °C, 5% CO_2_). Imaging was conducted by using 63X/1.2 WI C-Apochormat objective. Lamin-SS or Lamin-TM were excited with 514 nm laser line, pixel size was adjusted to 0.13 μm (zoom setting 4) and 256 x 256-pixel images were captured without averaging (195 ms scanning time per frame). In the FRAP experiment, images were collected with 250 ms intervals (249 images in total), and a bleaching was conducted after 9 scans. In the bleaching phase, a pre-drawn rectangular area of 75 x 10 pixels in the nuclear lamina was scanned 25 times (iterations) with 100% light intensity from 514 nm laser. The recovery was then followed for 240 frames.

### FRAP Data Analysis and Simulations

FRAP recovery curves were measured by using ImageJ FIJI-distribution.^50^ The drift of the nucleus during the imaging was corrected by using StackReg-plugin.^51^ Next the fluorescence was measured from the lamina and from the whole nucleus. The data was then normalized in Microsoft Excel for Mac (version 16.55) according to Phair & Misteli:^52^

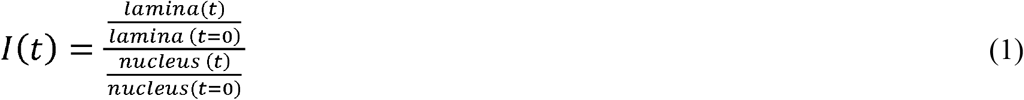

Where lamina(t) is fluorescence in the lamina at time point t, lamina(t=0) is fluorescence in the lamina before the bleach phase, nucleus(t) is the fluorescence of the whole nucleus at time point t and nucleus(t=0) is the fluorescence of the nucleus before the the bleach phase. Finally, the normalized recoveries were averaged.

Virtual Cell software^53,54^ was used to simulate the FRAP experiment and fluorescence recovery. The model contains a free Lamin-SS sensor which can bind to an immobile binding site in the lamina (single bound sensor), this binding can then lead into release of the sensor or tighter binding, simulating the situation where the sensor is engaged from both nanobodies (dual bound sensor). The release of the dual bound sensor was assumed to happen via single bound-state. The Lamin-TM sensor behavior was assumed to behave otherwise similarly, only the dual binding opportunity was missing. The reaction network schematic is visualized in Supplementary Figure 2. The Virtual Cell Models, “Lamin-SS_dual_binding” and “Lamin-TM_single_binding” by user “teihalai”, can be freely accessed within the VCell software (available at https://vcell.org).

### SensorFRET Imaging and Analysis

Live cells were seeded on glass-bottom slides coated with 20μg/mL fibronectin. DMEM was replaced with live cell imaging solution (cat #: A14291DJ, Thermo Fisher) supplemented with 10% FBS. Images were acquired using an inverted Zeiss LSM 710 (Oberkochen, Germany) confocal microscope using both 405 nm or 458 nm excitation wavelengths from an argon laser source. A 40x water immersion objective lens (NA = 1.1) was used for all imaging. Live cells were imaged in spectral mode using a 32-channel spectral META detector to record spectra of each pixel spanning wavelengths from 416 to 718 nm (with 9.7 nm spectral steps). Images were captured in 16-bit mode, scanned bi-directionally, and averaged 4 times. For sensorFRET based efficiency imaging, spectral images at both 405 and 458 nm excitation wavelengths were acquired. The normalized emission shape of the mTFP and mVenus fluorophores as well as the calibration parameter c (= 0.101) required for the sensorFRET analysis were experimentally determined from control cells expressing single fluorophores.^55^ Intensity images were further processed and analyzed using a custom Python code, which involves background subtraction and removal of saturated pixels. For each data set, the data was acquired for at least 5 images per condition per experiment. Images were masked manually on Fiji Image J.

### Paired FRET Measurements and Analysis

Ratiometric FRET imaging was used for FRET measurements involving paired FRET samples. Cell seeding and mounting was performed with similar protocol as in FRAP experiments. For live cell imaging, cells were placed in the microscope incubator (37 °C, 5% CO_2_). Zeiss LSM 780 laser scanning confocal microscope equipped with Plan Apochromat 63x/1.4 oil immersion objective was used for ratiometric FRET approach. FRET imaging and analysis was done by RiFRET method described previously.^56^ Briefly, the donor and acceptor were excited with a 458 nm line and a 514 nm line, respectively, from a multiline argon laser. The resulting fluorescence was acquired between 465–500 nm for donor emission and 535–650 nm for acceptor emission with a 32-channel QUASAR GaAsP PMT array detector. FRET channel emission was obtained with donor excitation (458 nm) and detected through the acceptor emission channel. Cells stably expressing either donor or acceptor probes alone was used to determine the spectral croTMalk. RiFRET plugin^12^ for ImageJ was used for croTMalk correction of each channel and to calculate pixel by pixel-based apparent FRET efficiency. The apparent FRET efficiency from individual cells prior to and after treatment was used for analysis.

### Fluorescence Lifetime Imaging Microscopy (FLIM)-FRET analysis

For FLIM, cells cultured in coverslips were fixed with 4% PFA for 10 mins, washed and stored in PBS at 4°C in dark before imaging. Prior imaging, the coverslips were mounted on imaging chamber and PBS was added to the chamber. Fluorescence lifetime imaging was performed using Leica STELLARIS FALCON confocal microscope equipped with Plan Apochromat 40x/1.25 motCORR glycerol immersion objective. Cells were excited with White Light Laser Stellaris 8 at 450 nm, and fluorescence lifetime times were recorded with HyD X detector, in the range 455 to 495 nm to obtain the photon arrival times specific to donor emission. The pixel-by-pixel photon arrival times were fitted for bi-exponential decay components using n-Exponential Reconvolution fitting model of Leica LAS X software to obtain mean lifetimes from individual cells.

### Statistical Analysis

Statistical significance was measured using an unpaired or paired, two-tailed Student′s t-test for data containing two groups. For data involving more than two groups, the Ordinary One-way Analysis of Variance (ANOVA) test was performed in order to obtain the statistical analysis for the data sets concerned. A further comparison of the groups was conducted using the Tukey (HSD) test so as to obtain significant differences between multiple groups. All statistical tests were conducted at a 5% significance level. GraphPad Prism was used for statistical analyses.

## Acknowledgments

We thank Jan Lammerding, Alice Varlet, Andrew Stephens, and Brenton Hoffman for thoughtful discussions and Heidi Peussa for help with the LMNA KO cells. The authors acknowledge the Biocenter Finland (BF), Tampere Imaging Facility (TIF), and the VCU Nanomaterials Characterization Core (NCC) for microscopy services. In addition, we wish to acknowledge Tampere University Virus Facility and Eric Dufour for the help in sgRNA design, Flow Cytometry Facility and Laura Kummola for the services and Light Microscopy Unit supported by HiLIFE and BE, Institute of Biotechnology, University of Helsinki, for the FLIM imaging. This project was funded in part by a National Science Foundation Graduate Research Fellowship (to B.E.D.), National Science Foundation awards CMMI 1653299 and CMMI 2135653 (to D.E.C.), National Institute of Health award R35 GM119617 (to D.E.C.), as well as Academy of Finland under the award numbers 308315, 314106, 33520 (to T.O.I.) and 332615 (to E.M).

