## Supplemental Figures for "Nuclear lamina strain states revealed by intermolecular force biosensor"

* Contributed equally

**Supplemental material:**

**Supplemental figures and figure legends:**

Supplemental Figure 1. Lamin-SS sensor distribution and apparent FRET efficiency in LMNA KO cells.

Supplemental Figure 2. Fluorescence recovery after photobleaching (FRAP) experiments of Lamin-SS and Lamin-SS-T binding to nuclear lamina.

Supplemental Figure 3. Schematic representation of the simulated FRAP experiment and reactions.

Supplemental Figure 4. The effect of actin cytoskeleton disruption on Lamin-SS FRET.

Supplemental Figure 5. Nuclear lamina organization after chromatin relaxation.

Supplemental Figure 6. Fluorescence recovery after photobleaching (FRAP) experiments of Lamin-SS after TSA treatment.

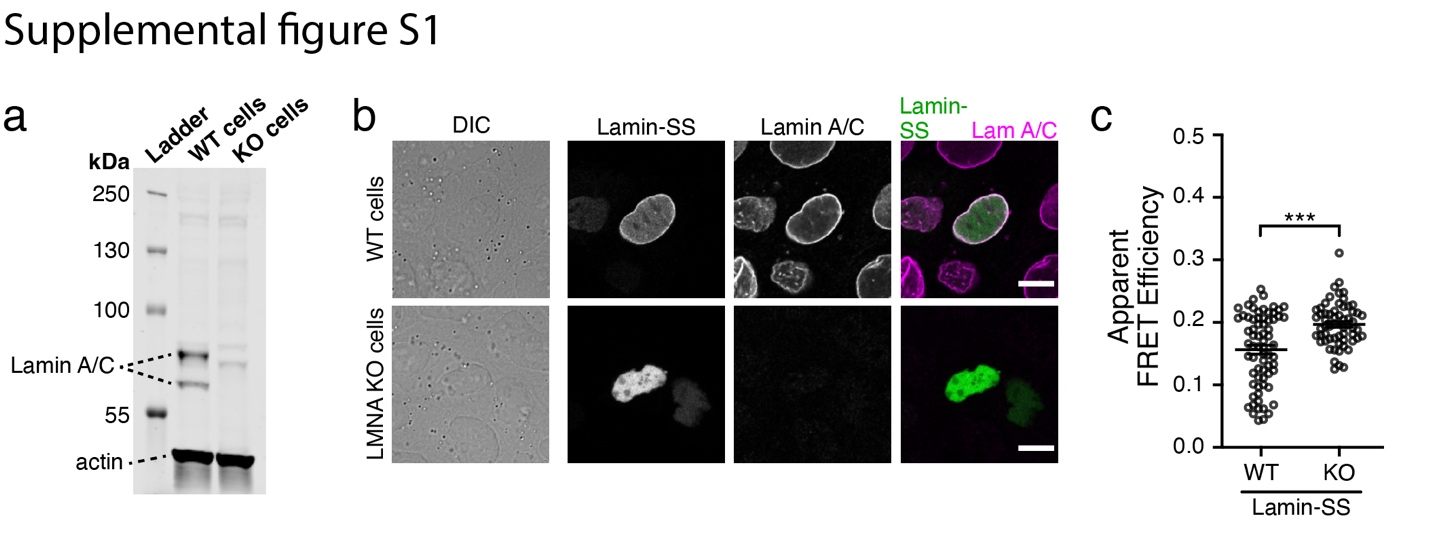

**Supplemental Figure 1. Lamin-SS sensor distribution and apparent FRET efficiency in LMNA KO cells.**

**a,** Immunoblot analysis of Lamin A/C in MDCK WT and LMNA KO cells showing successful knockout of LMNA gene in KO cells. **b,** Wild-type (WT) and LMNA KO cells transiently transfected with Lamin-SS. In LMNA KO cells Lamin-SS localization to nuclear lamina is lost. Scale bars 10 µm. **c,** Quantified apparent FRET efficiency (mean ± SEM) of Lamin-SS in WT and LMNA KO cells. Unpaired Student´s t-tests ((***) p ≤ 0.0001, t=4.65, df=126).

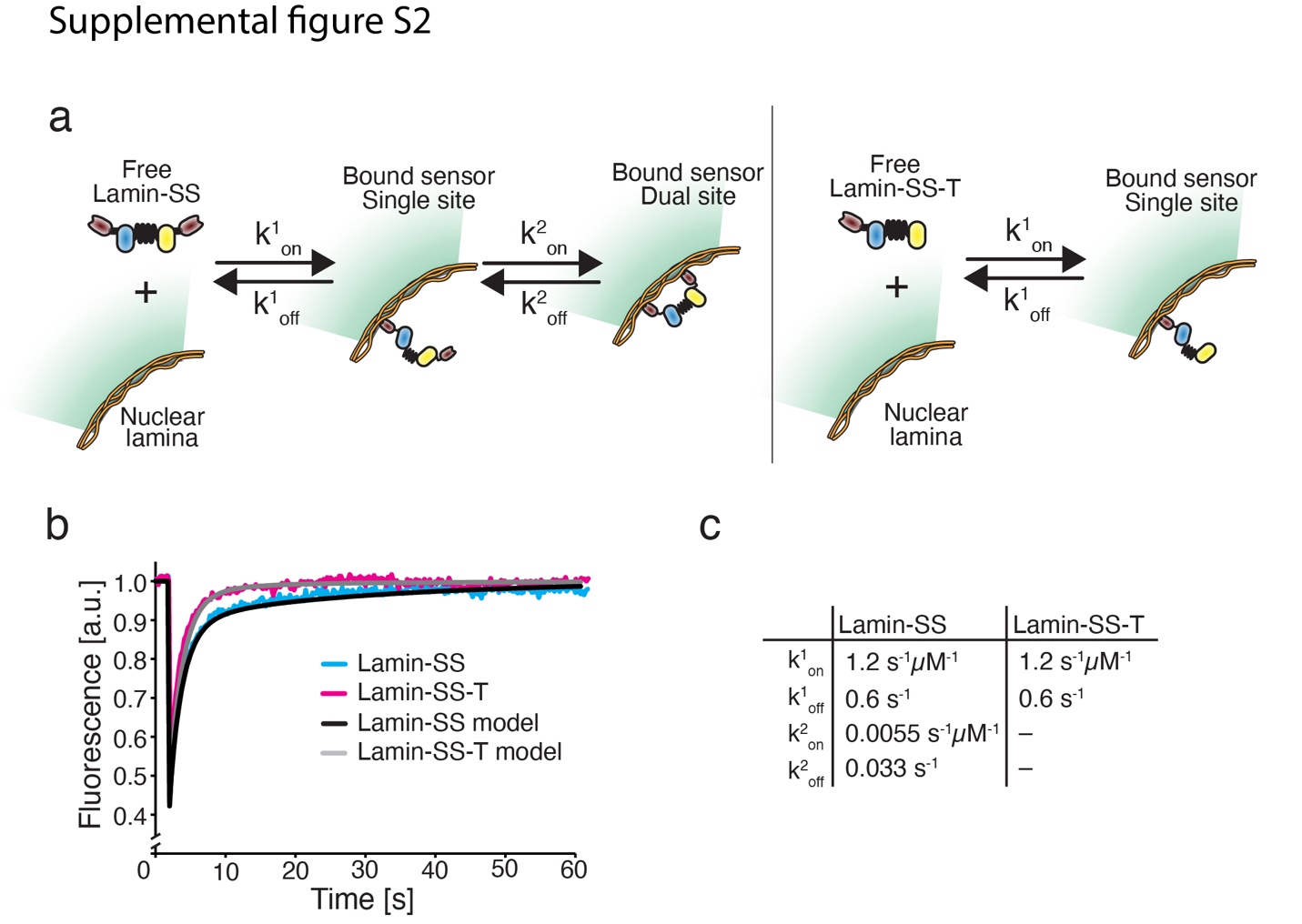

**Supplemental Figure 2. Fluorescence recovery after photobleaching (FRAP) experiments of Lamin-SS and Lamin-SS-T binding to nuclear lamina.**

**a,** Model of the Lamin-SS and Lamin-SS-T binding to nuclear lamina. Binding of Lamin-SS is assumed to proceed sequentially, from the binding of the first nanobody (single site) to the binding of the other (dual site). The release of the sensor from the lamina is modeled to proceed in reverse. Lamin-SS-T binding is limited to single site binding. **b,** Simulated recovery data together with the measured recoveries, see also main Figure 1. **c,** Binding pseudo on-rate and off-rate of Lamin-SS and Lamin-SS-T used in the simulations shown in b.

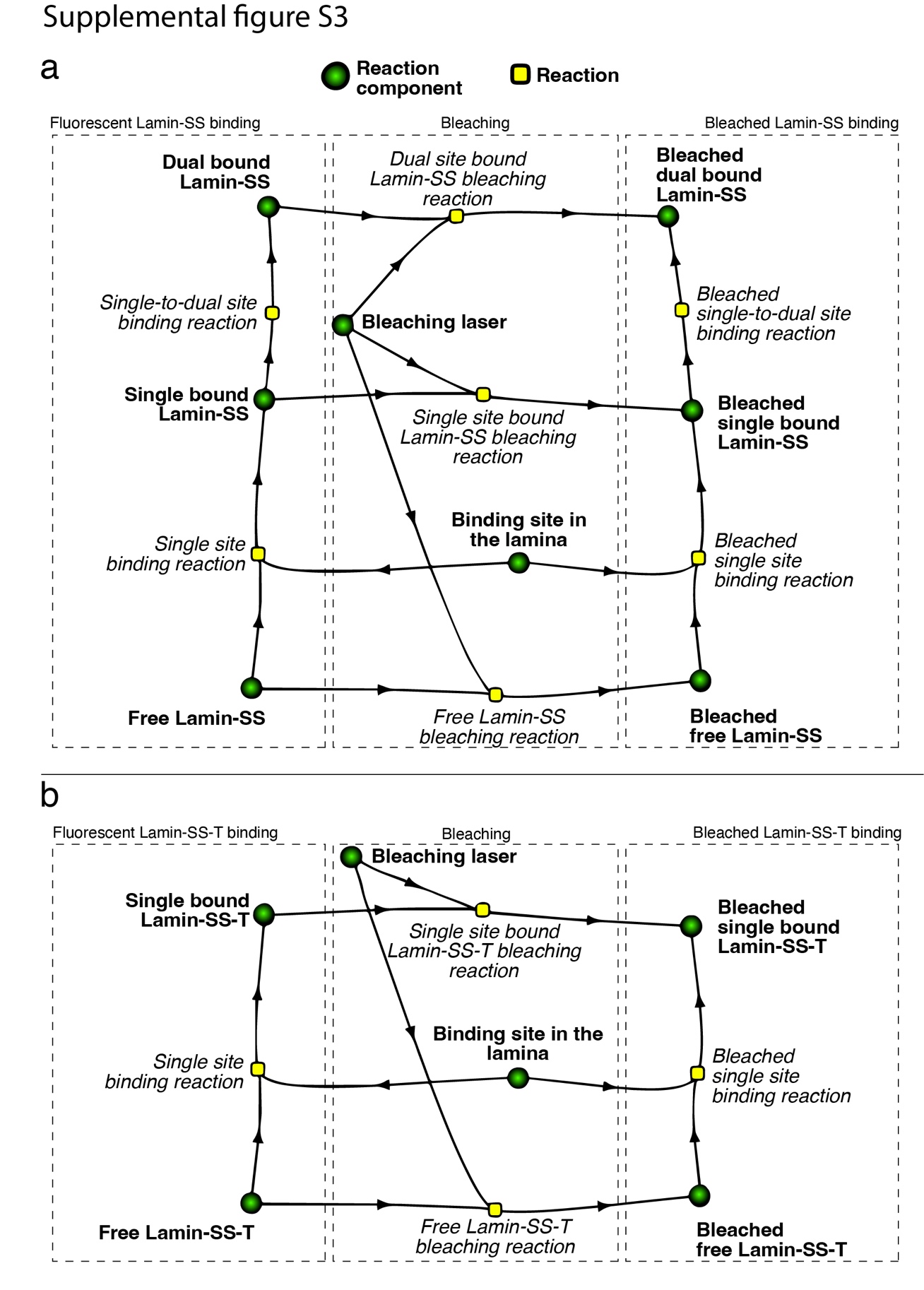

**Supplemental Figure 3. Schematic representation of the simulated FRAP experiment and reactions.**

**a,** Reactions of Lamins-SS during FRAP experiment. Freely diffusing Lamin-SS interacts with a binding site in the nuclear lamina (single site binding reaction) yielding Single bound Lamin-SS. This can further lead into Dual bound Lamin-SS (single-to-dual site binding reaction) or release of the sensor and binding site (reverse of single site binding reaction). Dual bound Lamin-SS can be released by reverse reaction leading into single bound Lamin-SS (reverse of single-to-dual site binding reaction). Bleaching is simulated by a local reaction between Bleaching laser, and free Lamin-SS, Single bound Lamin-SS and Dual bound Lamin-SS. The bleaching leads into appearance of bleached species of Lamin-SS. **b,** Reactions of Lamins-SS-T during FRAP experiment. Truncated sensor binding is limited to the reaction between free Lamin-SS-T and binding site in the lamina, yielding Single bound Lamin-SS-T (single site binding reaction). Similarly, as Lamin-SS, truncated Lamin-SS-T sensor is released, leading to Free Lamin-SS-T and Binding site in the lamina (reverse of Single site binding reaction). Fluorescent Lamin-SS-T molecules are bleached in the Bleaching reactions with Bleaching laser, yielding Bleached single bound Lamin-SS-T and Bleached free Lamin-SS-T.

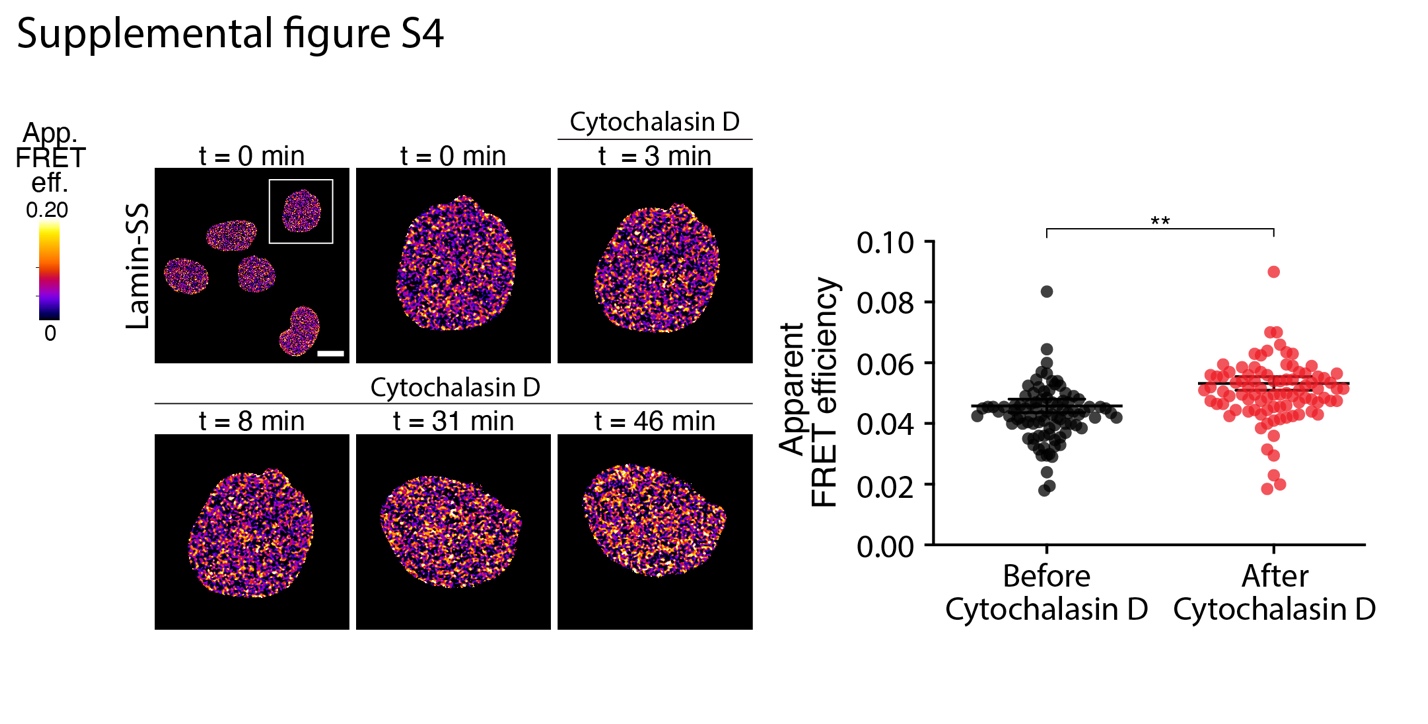

**Supplemental Figure 4. The effect of actin cytoskeleton disruption on Lamin-SS FRET**

Live cell imaging of Lamin-SS apparent FRET ratio during actin cytoskeleton disruption by Cytochalasin D (10 µg/mL). Scale bar 5 µm. Blow-up images of single nucleus indicate increase in the FRET (n= 92 cells, from 2 biological replicates). Paired Student´s t-test ((**) p<0.01, t=2.36, df=182).

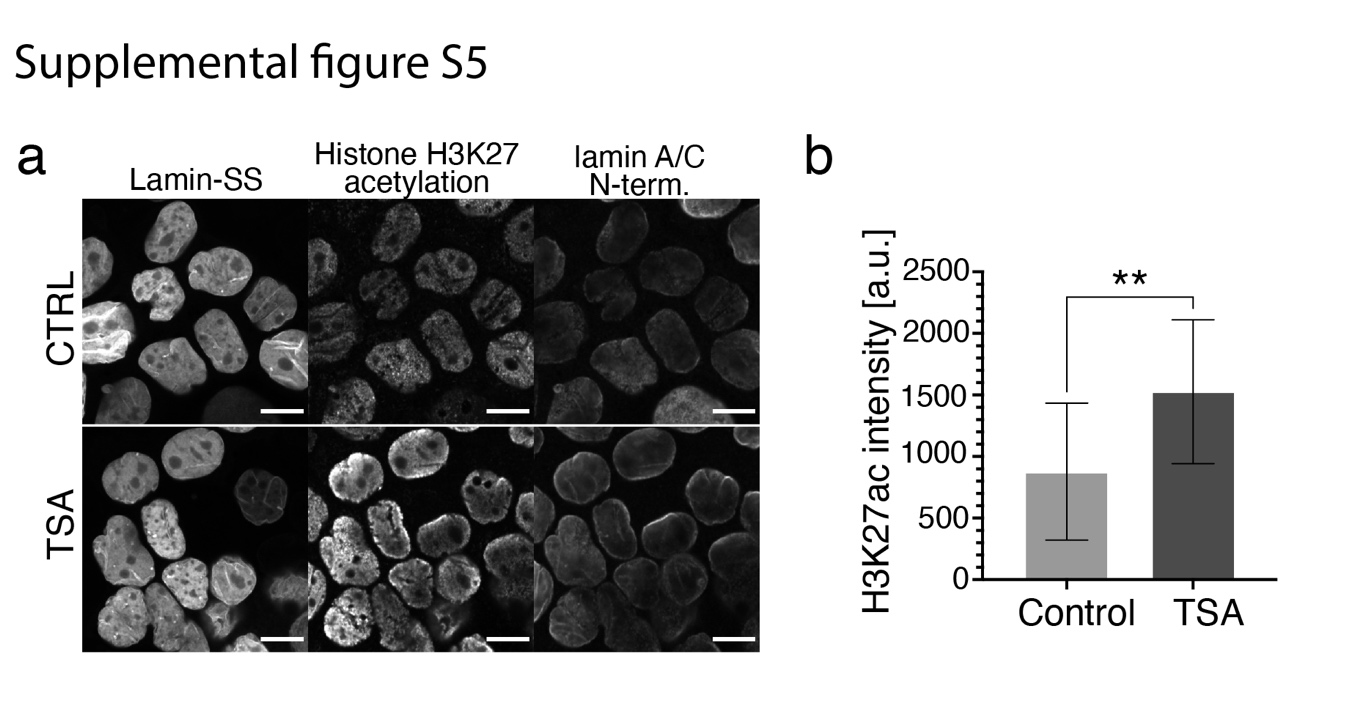

**Supplemental Figure 5. Nuclear lamina organization after chromatin relaxation**

**a,** Laser scanning confocal microscopy maximum intensity projection images of control (upper panels) and trichostatin A (TSA) -treated (600 nM, 4 h, lower panels) Lamin-SS expressing cells, immunolabeled against histone H3 lysine 27 acetylation (H3K27) and N-terminal part of A-type lamins. Scale bars, 10 µm. **b,** Quantification of nuclear fluorescence intensity of H3K27 acetylation labeling in control and TSA-treated cells (n=159 and n=151 cells, respectively, from 3 biological replicates). Unpaired Student´s t-test (** p=0.004, t=3.56, df=14).

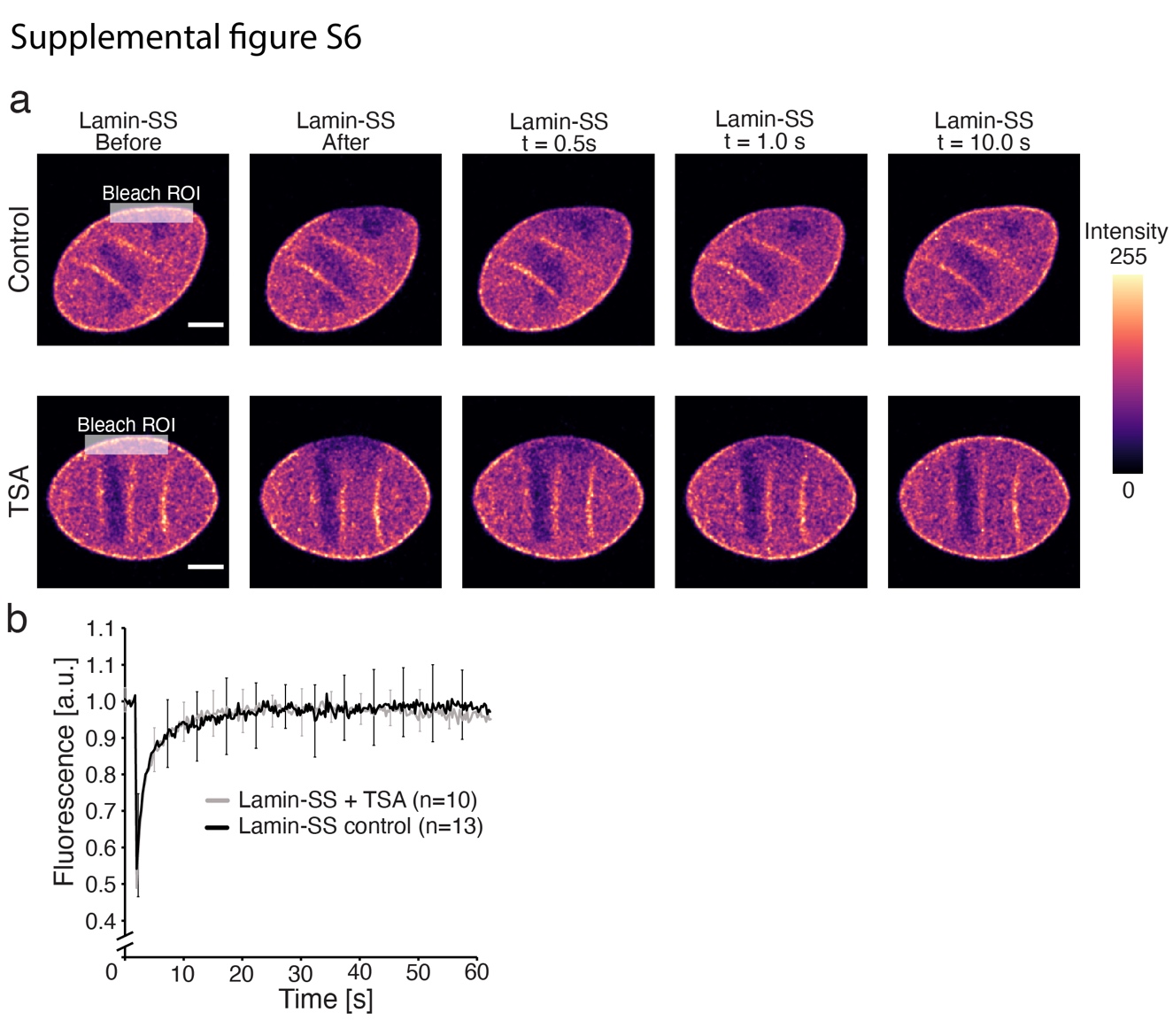

**Supplemental Figure 6. Fluorescence recovery after photobleaching (FRAP) experiments of Lamin-SS after TSA treatment.**

**a,** FRAP experiment with Lamin-SS expressing cells in control (upper panel) and after TSA treatment (600 nM, 4h). Bleached region of interest (ROI) is marked in the image. Scale bar 5 µm. **b,** Quantified and normalized fluorescence recoveries (mean ± standard deviation) of Lamin-SS in control and TSA treated cells (n=10 and n= 13, respectively, from 2 biological replicates).
